# SALVE: prediction of interorgan communication with transcriptome latent space representation

**DOI:** 10.1101/2025.08.12.669923

**Authors:** Jay Pavelka, Calvin Kin Voong, Peyton Schaal, Maggie P. Y. Lam, Edward Lau

## Abstract

Massive transcriptomics data allow gene relationships to be discovered from their correlated expression. We describe SALVE, a method to infer the associations between secretome-encoding transcripts and gene modules in a distal organ from RNA sequencing data. This method builds upon similar bioinformatics approaches by introducing transcriptome latent space representations and transfer learning to simultaneously increase discovery power and predict downstream functional associations. Applied to GTEx v8 data, we show the method readily recapitulates canonical endocrine relationships, including insulin and adiponectin signaling, while inferring new candidate organokines and their signaling modality. We also explore its utility for generating new hypotheses on cardiokine candidates and finding distal factors that may affect cardiac protein synthesis and metabolism. The predictions suggest a potential role of circulating galectin-3 (LGALS3) in regulating cardiac protein synthesis and homeostasis, which can be recapitulated in part in human induced pluripotent stem cell (hiPSC)-derived cardiomyocytes. This method may aid in ongoing efforts to delineate interorgan communications and endocrine networks in various areas of study.

**New & Noteworthy:** We describe a bioinformatics strategy to find associations between the secretome-coding genes and functional pathways of two organs. This approach may be used to find crosstalk signals between the heart, adipose, liver, and other tissues. Applied to GTEx data, it suggests a potential role of circulating galectin-3 in regulating cardiac protein synthesis and homeostasis pathways.

## Introduction

Organismal homeostasis requires the coordinated function of organ systems. For instance, the adipose tissue has emerged as an endocrine hub with the ability to control whole-body energy homeostasis through the release of organokine signals (1). In parallel, the risks of many age-associated diseases such as heart failure are significantly modified by systemic factors including diabetes and metabolic syndromes. Interorgan regulations have thus garnered significant attention both as a way to understand disease mechanisms and also as prime targets for therapeutic manipulation (2). Nevertheless, important knowledge gaps remain unfilled regarding tissue crosstalk and how it goes awry in diseases, and in general, the full gamut of organokine proteins that act as messengers of interorgan communication remains poorly mapped.

The human genome contains almost 3,000 genes that encode secreted proteins, accounting for ∼15% of the protein-coding genome (3). Current knowledge of endocrine messengers that mediate interorgan signaling networks however remains vastly incomplete. Targeted dissections of cross-tissue communication signals require a priori hypotheses on the specific tissues they regulate and thus remain challenging without unbiased discovery. Advances in proteomics technologies have allowed circulating molecules to be profiled in depths. However, with the plasma proteomes being a combined pool of effluents of every tissue, it remains difficult to disentangle which tissue an identified plasma protein may originate from, which is further complicated by the fact that many plasma proteins have multiple simultaneous tissue sources (4). Advances in proximity labeling of tissue-specific secretomes have now enabled the unbiased discovery of the tissue origins of circulating proteins (5–7), but it remains technically challenging to decipher which tissues they then travel to and the biological messages they may carry (8, 9).

An emerging powerful alternative is to infer interorgan associations directly from large-scale transcriptomics data. Co-expression analysis is a foundational bioinformatics strategy for discovering functionally related genes within a cell type or tissue (10, 11), by assuming a latent regulatory architecture that co-modulates functionally-related genes in a manner such that they covary across a panel of naturally variable samples. Several works in the past decade have generalized co-expression analysis toward two or more tissue transcriptomes, demonstrating it has the potential to find co-regulated genes across tissues to infer interorgan relationships (12–16). For example, an analysis by the GTEx Consortium found *DPP4* as a heart-to-blood communication factor through their cross-tissue co-variance (15). Subsequently, Seldin, Lusis, and colleagues performed co-expression analysis on five organs in the Hybrid Mouse Diversity Panel (HMDP) study to discover endocrine interactions (14). This paper introduced a rank-based metric, S_sec_, to further prioritize bona-fide signals, calculated as the sum of the negative natural logs of correlation P values between each source-tissue secreted protein-coding transcript across all target-tissue transcripts. Outliers of S_sec_, that is, source-tissue genes with unusually high correlations with target tissue genes, are then nominated for further filtering. This method led to the discovery of new endocrine factors and has since been applied to additional studies (13, 17). Other studies likewise demonstrated it is possible to use population variance in large transcriptomes data to find bona-fide endocrine signals, that can then be experimentally validated in vivo (12, 16). Despite the many insights from these studies, there remains room for continued method improvements. For example, current methods may bias toward genes that interact modestly with many targets even if they are not relevant to crosstalk pathways of interest. Secondly, the downstream functional pathways of a signaling event are often not directly tested as correlates but inferred post-hoc via overrepresentation analysis of top genes. Third, many gene-to-gene relationships need to be tested, which can incur false positives under low sample sizes.

Here, we explore a complementary approach to predict endocrine signals, which considers **<u>s</u>**ecretome **<u>a</u>**ssociations with transcriptome **l**atent **<u>v</u>**ariables between tissu**<u>e</u>**s (SALVE). This approach extends on prior efforts by simultaneously inferring a source-tissue organokine and its correlated downstream function in target tissues via gene-to-module co-expression. To infer gene module expression, we use the pathway-level information extractor (PLIER) algorithm to create low-dimension representations of target tissue transcriptomes that map to functional annotations. The resulting predictions recapitulate known cross-tissue signaling modalities while generating new hypotheses of novel crosstalk protein candidates. Applying the approach to GTEx v8 human transcriptome data, we highlight a predicted role of galectin-3 and thrombopoietin on protein synthesis pathways in the heart, which are recapitulated experimentally in hiPSC-derived cardiomyocytes. The presented approach may be applicable to predicting tissue crosstalk in multiple organ systems using different data sets.

## Materials and Methods

### Data retrieval and normalization

Human RNA-sequencing read counts from the Common Funds Genotype-Tissue Expression (GTEx) v8 study (18) containing ∼17,382 transcriptomes from ∼948 donors (dbGaP phs000424.v9.p2.c1) were retrieved the GTEx data portal (https://www.gtexportal.org). The read counts were first normalized using the Variance Stabilizing Transformation wrapper from the R DESeq2 package (19). Batch effects were corrected for using ComBat with the sva R package (20), sequentially correcting extraction batch (SMNABTCH), sequencing batch (SMGEBTCH), tissue ischemic time (SMTSISCH) in 300 min intervals (0, 300, 600, 900, 1200, >1200), death types (DTHHRDY), and decadal age bins (**Supplemental Figure S1**).

### PLIER Model training

Human Protein Atlas (HPA) annotations of protein localization (3) were retrieved using the hpar R package (21) to categorize transcripts into “Secreted to blood” or otherwise. Transcripts in the latter were used for PLIER model training. (22). Briefly, PLIER is a matrix decomposition algorithm that produces low-dimension representations (latent variables) of a transcriptome that map to known pathways given prior knowledge annotations. As described in Mao et al. (22), the PLIER algorithm minimizes the following terms:

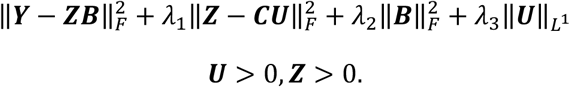

This approach minimizes the reconstruction error of the original gene expression matrix Y into the loading sand latent variable matrices ***Z*** and ***B***. This dimension reduction is constrained by mapping to the prior knowledge matrix ***C***, while rewarding sparsity of ***B*** and ***U*** through the penalty terms. We used function provided by PLIER and the multiPLIER scripts to determine free parameters and minimize reconstruction error. Subsequently, the transcripts were filtered for samples that contains read counts from both the donor (source) and acceptor (target) tissue for each source-target pair. Near zero variance genes from the target tissue were first removed using a function within the caret R package (23). Following the approach described in Taroni et al., the single ***Z*** matrix learned from the full GTEx data, were transferred onto the target tissue gene matrix to generate a new ***B*** matrix, using the transfer learning wrapper function within the multiPLIER R package (24), as follows:

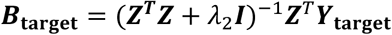

### Correlation matrix and protein ranking calculation

For each source-target tissue pair, a Pearson’s correlation matrix was generated across subjects between all source-tissue HPA secretome genes and the target tissue ***B*** matrix, using a correlation function within the Hmisc R package 8/12/25 11:49:00 AM. Family-wise error rate was controlled using Bonferroni correction based on the number of correlations in each source-target tissue pair. A Bonferroni P value (adj.P) of 0.005 or below is considered significant.

The S score for source-tissue gene rank prioritization for each source-target tissue pair *i*, *j*; *i* ≠ *j* was calculated from the Pearson’s correlation matrix by taking the sum of all negative logs of all Bonferroni corrected P values between the source tissue gene *g* for all target-tissue latent variables *l*. To limit outsized contributions by a single latent variable association, P is capped at 1e–6 from below:

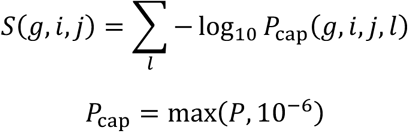

The standardized S* score for a source tissue gene in each source-tissue pair is then calculated analogous to a Z score:

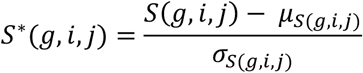

### Human induced pluripotent stem cell culturing and differentiation

Human induced pluripotent stem cells (hiPSCs) were sourced from the Allen Institute for Cell Science (AICS-52, expressing mEGFP-tagged MLC-2a on the WTC-11 parent line). For culturing, cells were thawed and plated onto Geltrex (Gibco #A1413301) coated 6-well plates and expanded using mTeSR Plus media (StemCell #100-0276). Y-27632 ROCK inhibitor (SelleckChem) was applied for 24 hr after initial seeding. Cells were then expanded and passaged to 12-well plates with a fixed density cell count of 0.3 million cells per well to start differentiation. Once the cells become 80% confluent (day 0), media was switched to RPMI-1640 containing glucose (Gibco CAT#11875093) supplemented with B27 without insulin (Gibco CAT#17504044) with 6 µM CHIR-99021 (SelleckChem). On day 2 of differentiation, CHIR-99021 was removed from the media and on day 3, IWR-1-endo (SelleckChem) was introduced as a Wnt inhibitor at 5 µM (SelleckChem). On day 5 of differentiation, IWR-1-endo was removed via media change with RPMI-1640 and B27 without insulin. On day 7, the media was replaced with RPMI-1640 with glucose and supplemented with B27 with insulin. On day 11, media was changed to RPMI-1640 without glucose (Gibco #11879020) and supplemented with B27 with insulin to select for differentiated cardiomyocytes. After contractile cells were observed on day 13 of differentiation, cells were passed and plated for experiments. Cells were stained for F-actin using the CellMask Orange Actin Tracking Stain (ThermoFisher #A57244) and ProLong Diamond Antifade Mountant with DAPI (ThermoFisher #P36962) alongside MLC-2a-mEFGP to verify differentiation.

### Organokine treatment and RNA extraction

Recombinant human galectin-3 protein (abcam #ab50236) was purchased from Abcam and was reconstituted in 0.1% BSA to a stock concentration of 100 µg/mL and used as a final concentration of 14 ng/mL, in line with clinical range (25–27). Recombinant human thrombopoietin protein (ThermoFisher #PHC9511) was reconstituted in 0.1% bovine serum albumin to a stock concentration of 2 µg/mL and used as a final concentration of 0.1 ng/mL (28). Cells were treated with the final concentrations of adipokines for 48 hr and then harvested by scraping in TRIzol reagent. RNA was extracted using Direct-zol RNA mini prep kits (Zymo Research #R2050). RNA yield and integrity was assessed using Qubit assays prior to sequencing (Thermo).

### RNA sequencing and analysis

RNA sequencing was performed on Illumina short-read polyA+ libraries for 40M 150-nt paired-end reads per sample (Novogene). RNA sequencing reads were aligned to GENCODE v44 reference genome and annotations using STAR v.2.7.11a (29) with arguments --sjdbOverhang 149 --outSAMtype BAM SortedByCoordinate. Transcript assembly and quantification was performed using StringTie v.2.1.1 (30). The resulting gene count matrix was analyzed using the DESeq2 package in R (19). Genes with total counts ≥ 64 across samples were retained. Differential expression results with DESeq2 apeglm lfcShrink s-value ≤ 0.1 are considered significant. Gene set enrichment analysis (GSEA) was performed using the fgsea package in R against MSigDB C2 annotations. Enrichment results with FDR-adjusted P ≤ 0.1 are considered significant.

## Results

### Transfer learning and latent space co-expression predicts cross-tissue signals

We reason that a dimension reduction approach will allow us to find functionally relevant latent variables in a target tissue, and at the same time reduce multiple testing burden by performing co-expression analysis only between source tissue secreted genes with a limited number of target tissue latent variables. PLIER (Pathway Level Information Extractor) (22) is a context-aware dimension reduction algorithm that favors decomposition of gene expression matrices into a gene-by-latent-variable matrix Z and a latent-variable-by-sample matrix Z that align sparsely with one or few known pathways or gene ontology terms. Subsequently, Greene and colleagues further show that the PLIER loading matrix Z may be pre-trained from other data than the ones being analyzed using the large recount2 data collection (∼37,000 transcriptomes) in a transfer learning paradigm (24). This creates a generalizable latent space that represents gene expression architecture in a tissue upon which new data can be projected. In prior work, we showed that the GTEx v8 data set (∼17,000 transcriptomes) can be used to train a large Z matrix to aid in data interpretation of smaller-sample-size transcriptomics data (31). We therefore decided to use PLIER to extract latent space representation of transcriptome profiles prior to interorgan co-expression analysis, which should allow us to test the correlation of secretome-coding genes with target tissue gene modules that map to interpretable pathways, and thus focus on only endocrine signaling that affects the biology of interest in a target tissue.

We retrieved GTEx v8 RNA-sequencing read counts data, comprising 54,592 transcripts and 17,382 samples across 54 tissue subtypes, with an average of 322 samples per tissue within the data set (**Figure 1A Step 1**). However, when considering a pair of tissues, the sample size decreases to a median of 96 shared samples, illustrating the data size problem. Following normalization and batch correction, we divided the data set into secretome and non-secretome coding transcripts (**Figure 1A Step 2–4**) and trained a model of underlying gene expression structure (i.e., the PLIER latent variables) using the entire GTEx v8 collection (∼17,000 transcriptomes) (**Figure 1A Step 5**). This trained model is then applied to the smaller scale cross tissue data in a transfer learning paradigm (**Figure 1A Step 6–7**), and finally calculating the correlation coefficients of all pairwise source-tissue relationships between source-tissue secretome transcripts and target-tissue transcriptome latent variables, followed by stringent multiple testing correction (**Figure 1A Step 8**). Finally, we used a rank-based metric (S*) to sort organokines from a known source organ based on the total number of latent variable correlations it has with any target organs (see Method), which is then to yield a compendium of prioritized endocrine signaling candidates with S* score ≥ +2 within a source-target tissue pair. In total, we nominated ∼1.3 million significant correlations across GTEx tissues involving 602 unique organokines and 696 transcriptome latent variables in 54 GTEx tissues (**Figure 1 Step 9**).

**Figure1:**
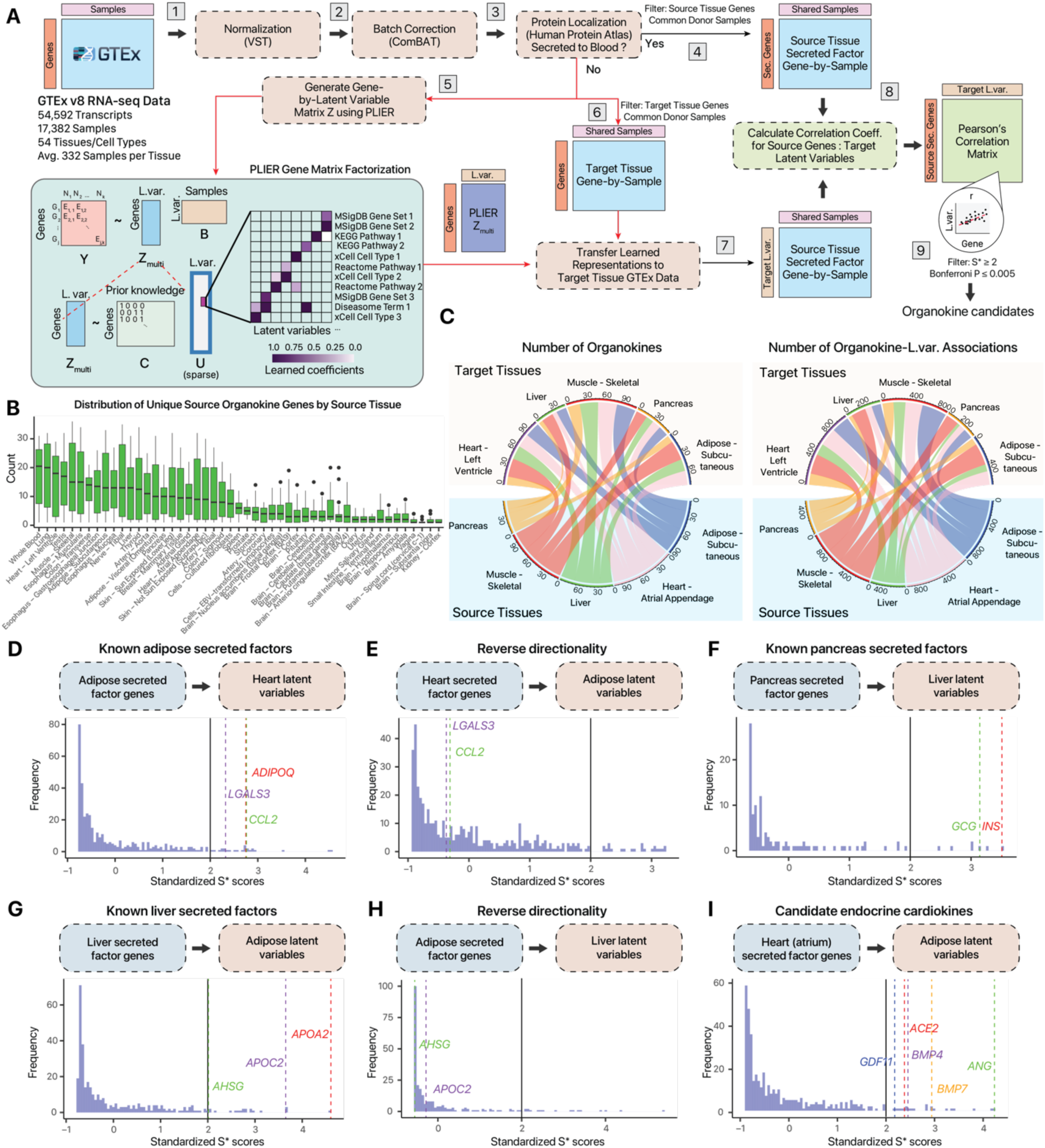
Inferring endocrine relationship via Secretome Association with Latent Variables between Tissues (SALVE) **A.** Presented workflow. Large transcriptome data from GTEx v8 is (Step 1) normalized and (Step 2) sequentially batch corrected. Transcripts are then categorized based on whether the protein products are known to be secreted to blood in the Human Protein Atlas (Step 3). Secreted factor genes from the source tissues of interest are extracted (Step 4). All non-secreted factor genes in the entire GTEx data set are then used to train a PLIER Z matrix that relates genes to latent variables mapping to known functional annotations or pathways (Step 5). This learned matrix is used to create a low-dimension representation of gene expression in the target tissue of interest (Step 6 and 7). A correlation matrix between all source tissue secreted factor genes and target tissue latent variables is created for each source-target tissue pair (Step 8). **B.** Boxplot showing distribution of significant correlation between each source tissue (x-axis) with all target tissues. **C.** Chord diagram showing the number of significant organkines (left) and organokine-latent variable correlations (right) (adj.P ≤ 0.01, S* ≥ 2) among prioritized organokines across several tissues of interest. **D.** Histogram showing the Standardized S* score of associations between adipose (subcutaneous) secreted factor coding transcripts and heart (left ventricle) latent variables. Organokines with Standardized S* scores ≥ 2 (black line) are prioritized. Known adipokines including *ADIPOQ*, *LGALS3*, and *CCL2* that are captured in the analysis are highlighted **E.** As in D, but showing heart secreted factors to adipose (subcutaneous) latent variables associations. Adipokines no longer have significant scores. **F.** As in D, but showing pancreas secreted factors to liver latent variables associations, highlighting known pancreas secreted hormones glucagon and insulin. **G.** As in D, but showing liver to adipose (subcutaneous) latent variables associations, highlighting the known hepatokine *AHSG*. Apolipoproteins *APOA2* and *APOC2* also show significant scores. **H.** As in G, but showing adipose to liver associations. **I.** As in G, but showing heart (atrial appendage) to adipose associations, highlighting candidate cardiokines that may participate in endocrine signaling including *GDF11*, *ACE2*, *BMP4*, *BMP7*, and *ANG*.

Across all pairwise relationships among the 54 tissues examined, whole blood had the greatest number of significant cross-tissue correlations as a source tissue, reflecting its central role in tissue crosstalk in the body (**Figure 1B**). Although not all GTEx tissues (e.g., skin) might be expected to participate extensively in endocrine signaling, we retain all pairwise tissue analysis here to present a systems-wide resource (**Supplemental Table S1**). Clustering analysis shows that organ pairs with the greatest numbers of mutual significant associations are centered on muscles, adipose, pancreas, the heart, and the lungs alongside traditional endocrine tissues like the thyroid, pancreas, and pituitary gland, suggesting the organokines secreted from these tissues are likely to influence the gene expression profiles of each other (**Supplemental Figure S2**). In contrast, brain tissues form a separate cluster that preferentially associate with one another, whereas the ovary, kidney cortex, uterus, and the minor salivary glands have few associations, suggesting these tissues are less likely to affect or be affected by other tissues via organokine genes. Based on this result, we focused on several active secretory organs that are important to cardiometabolic regulations, including the heart, skeletal muscle, and adipose, which form a network of interorgan connections with each other (**Figure 3C**).

We assessed whether the approach recapitulates known endocrine proteins and the tissues they signal to. Focusing on specific source/target tissue pairs, we assess the S* scores of candidate adipokines between adipose tissue and the heart, showing that hemokine (C-C motif) ligand 2 (CCL2/MCP-1) and adiponectin (*ADIPOQ*) scored highly (S* ≥ 2) (**Figure 1D**); implying the adipose secreted pools of these organokines are associated broadly with the gene expression landscape of the heart (left ventricle) tissue. This co-expression pattern is no longer observed when the directionality of the tissue-pairs is reversed (heart (left ventricle) ∼ adipose (subcutaneous) (**Figure 1E**). Other examples highlight insulin (*INS*) and glucagon (*GCG*) as positive outliers among pancreas secreted factor genes that associate with liver latent variables (**Figure 1F**), and fetuin A (*AHSG*) as an outlier in the liver (**Figure 1G**). The scoring likewise dissipated when the source and target tissues are swapped; hence, the nominated organokines influence target tissue gene expression landscape specifically and unidirectionally (**Figure 1H**).

A potential utility of this analysis is to uncover endocrine candidates originating from various tissues. For instance, emerging studies have highlighted a large repertoire of extracellular proteins that are secreted by cells in the heart, or cardiokines (32, 33). Beyond the classic examples of natriuretic peptides however, which of the discovered cardiokines function in endocrine as opposed to paracrine fashion remains a subject of research. An example query of heart (atrial appendage) to adipose (visceral - omentum) correlations uncovered several candidates, including *BMP4*, *BMP7*, and *GDF11*, hinting at their involvement in long-range signaling from the heart (**Figure 1I**). Therefore, the analysis is able to recapitulate known canonical endocrine signals while generating new hypotheses on endocrine candidates.

### Latent space co-expression captures downstream effects of organokine signals

We next explored the compositions of significant gene modules for a particular secretome gene to assess whether our approach recapitulates known biology. From the adipokines along the adipose to heart signaling axis above, we focus on the significant secretome to gene module relationships (S* ≥ 2, adj.P ≤ 0.05) that involve adiponectin. Adiponectin is an extensively characterized adipokine known to modulate immunity and inflammation, calcium signaling, and insulin sensitivity in multiple organs including the heart (1, 34). The predicted heart transcriptome latent variables that are most correlated with adiponectin in the adipose are consistent with these known roles, as evidenced by the high correlations of adipose ADIPOQ with latent variables that map to metabolism, immunity and calcium signaling pathways at adj.P ≤ 0.005, including L.var.57 (REACTOME Innate Immune System) and L.var.509 (KEGG Calcium Signaling Pathway (**Figure 2A**). A closer inspection of the transcriptome representations in each individual GTEx sample confirms a robust and significant correlation between the read counts from adiponectin against the normalized levels of the latent variables (**Figure 2B–C**).

**Figure 2:**
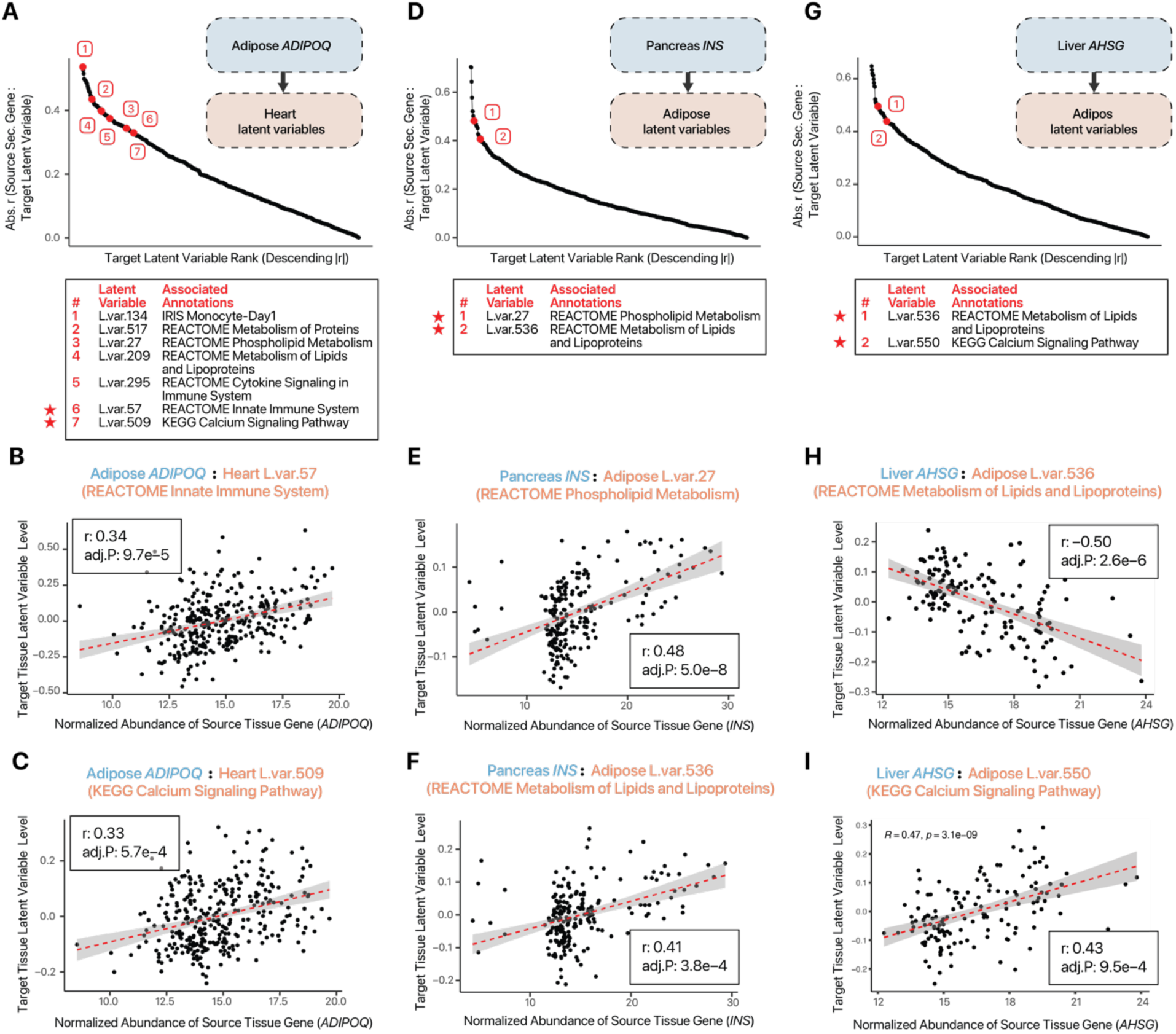
Associations between canonical endocrine signals and downstream pathway.s. **A.** Dot plot showing the absolute correlation of heart (left ventricle) latent variables with adiponectin (*ADIPOQ*) transcript level in the adipose tissue (subcutaneous). Selected significant (adj.P ≤ 0.005) latent variables (L.var.) are numbered, in red. Latent variables are ranked by descending absolute Pearson’s correlation coefficient (r). **B.** Scatter plot showing the correlated expression (r: 0.34, adj.P: 9.7e–5) of adipose *ADIPOQ* and heart (left ventricle) latent variable 57, which maps to the REACTOME Innate Immune System term. Each data point is an individual donor sample with both adipose and heart transcriptomes in GTEx v8. Red dashed line: best fit linear fit. Ribbon: s.e. **C.** As in B, but for the association between adipose *ADIPOQ* and heart latent variable 509 (KEGG Calcium Signaling Pathway) (r: 0.33, adj.P: 5.7e–4) **D.** As in A, but for adipose (visceral/omentum) latent variables associated with insulin (INS) transcript levels in the pancreas. **E.** As in B, but for the association between pancreas INS and adipose latent variable 27 (REACTOME Phospholipid Metabolism) (r: 0.48, adj.P: 5.0e–8) **F.** As in B, but for the association between pancreas INS and adipose latent variable 536 (REACTOME Metabolism of Lipids and Lipoproteins) (r: 0.41, adj.P: 3.8e–4) **G.** As in A, but for adipose (subcutaneous) latent variables associated with fetuin-A (AHSG) transcript levels in the liver. **H.** As in B, but for the association between liver AHSG and adipose latent variable 536 (REACTOME Metabolism of Lipids and Lipoproteins) (r: –0.50, adj.P: 2.6e–6) **I.** As in B, but for the association between liver AHSG and adipose latent variable 550 (KEGG Calcium Signaling Pathway) (r: 0.43, adj.P: 9.5e–4)

In parallel, pancreas insulin (*INS*) is correlated with adipose tissue L.var.27 (REACTOME Phospholipid Metabolism) and L.var.536 (REACTOME Metabolism of Lipids and Lipoproteins), consistent with the well characterized effects of insulin on adipose tissues (**Figure 2D–F**). Similarly, fetuin-A (*AHSG*), an abundantly secreted hepatokine, is associated with latent variables that reflect its known signaling modality. Fetuin A is known to bind multiple cell-surface receptors to regulate biological processes from bone remodeling to protein metabolism, and insulin signaling; and is known to suppress lipid metabolism by down-regulating CD36 (35). Along the liver to heart (left ventricle) axis, we find that liver *AHSG* is correlated with cardiac lipid/lipoprotein metabolism and calcium signaling latent variables (**Figure 2G**). Individual examination of the correlation plots shows a negative correlation with L.var.536 (REACTOME Metabolism of Lipids and Lipoproteins) and positive correlation with L.var.550 (KEGG Calcium Signaling Pathway) (**Figures 2F–H**). The decrease in these pathways agrees with current observations that suggest fetuin-A decreases in incidences of cellular apoptosis, calcium metabolism, and lipid metabolism (35). Finally, our results are consistent with the endocrine role of adipose lipopolysaccharide binding protein (*LBP*) (**Supplemental Figure S2**), previously shown to be an adipokine candidate in mice and humans that signals to the liver to modulate glucose and lipid metabolism (12). Therefore, SALVE predictions recapitulate prior established endocrine signaling events.

### SALVE nominates candidate endocrine modulators of biological pathways of interest

We next examined a use case of SALVE to generate hypotheses on organokine candidates associated with a biological process of interest. Cardiac protein synthesis and protein metabolism are critically associated with cardiac hypertrophy, atrophy, and other physiological adaptations. We therefore explored the secreted factor genes from cardiometabolically relevant organs that are significantly correlated with the ribosome associated latent variable in the heart as a proxy of protein synthesis. We find multiple source tissue genes predicted to be associated with the cardiac ribosome term at S* ≥ 2 and adj.P ≤ 0.005, primarily in skeletal muscle, adipose, and the liver (**Figure 3A**). The predictions nominated several genes coding for known and potential adipokines (*ADIPOQ*, *CCL2*/*MCP-1*, *IGFBP2* and *LGALS3*), hepatokines (*AGT*, *THPO)* and myokines (*IL15*). By contrast, distinct tissues and organokine genes predominate upon querying associations with lipid metabolism (**Figure 3C–D**) and collagen formation (**Supplemental Figure S4**) in the heart. Individual inspections of associations confirm significant correlation between normalized source tissue expression and target tissue latent variable level across individual donors (**Figure 3E**).

**Figure 3.**
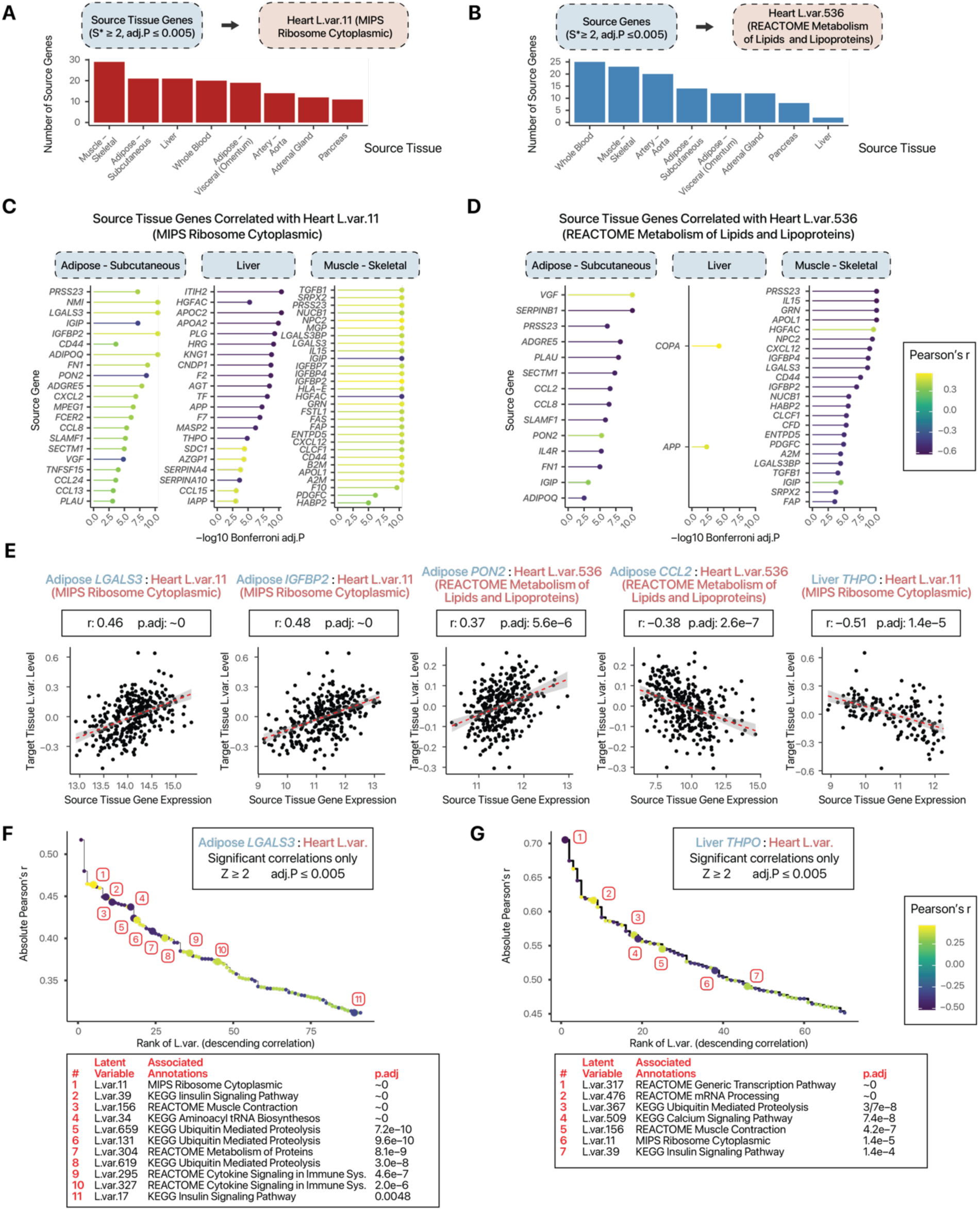
Candidate endocrine modulators of cardiac protein synthesis and lipid metabolism. **A.** Bar chart showing the number of significant organokines from several cardiometabolically relevant tissues that are significantly associated with heart latent variable 11 with MIPS Ribosome Cytoplasmic annotation. **B.** As in A, but for organokines associated with heart latent variable 536 (annotated with REACTOME Metabolism of Lipid and Lipoproteins). **C.** Dot plot showing the correlation coefficient and significance of organokines from the adipose tissue (subcutaneous), liver, and skeletal muscle with heart latent variable 11 (MIPS Ribosome Cytoplasmic). **D.** As in C, but for organokines associated with heart latent variable 536 (REACTOME Metabolism of Lipid and Lipoproteins). **E.** Scatterplot showing the correlated expression of adipose and liver organokines with heart ribosome and lipid metabolism latent variables. Each point is data from an individual donor. Red dashed line: best fit linear fit. Ribbon: s.e. **F.** Dot plot showing the absolute correlation of adipose (subcutaneous) LGALS3 with heart (left ventricle) latent variables. Only latent variables mappable to prior knowledge and with Bonferroni P ≤ 0.005 are shown. Color: Pearson’s correlation coefficient. Several latent variables of interest including those associated with protein synthesis and degradation pathways are highlighted. **G.** As in F, but for correlations between liver THPO with heart (left ventricle) latent variables.

To further examine the functional contexts of these organokines, we focus on galectin-3, a member of the galactin family encoded by the *LGALS3* gene and that binds β-galactosides. In the clinical setting, galectin-3 has been of growing interest due to its potential as a prognostic and diagnostic biomarker for heart diseases including heart failure (36). In our predictions, the two most significant signals of galectin-3 with cardiac ribosomes originate from the adipose and the liver, two organs important in cardiometabolic diseases. Among the most significant co-expression correlations, galectin-3 shows a strong influence on processes involving ribosomes, ubiquitin mediated proteolysis, immune responses, and the insulin signaling pathway (**Figure 3F**). As a second example, we consider thrombopoietin (*THPO*), a known hepatokine whose expression in the liver is correlated with ribosome annotated latent variable in the heart (**Figure 3C**). Among the top correlations, thrombopoietin is predicted to have a significant influence on ribosome as well as diverse processes including mRNA processing, ubiquitin-mediated proteolysis, and calcium signaling (**Figure 3G**). Taken together, SALVE implicates complex influences of these circulating factors on the heart and generates specific hypotheses on functional impact, which may be tested in various experimental models.

### Validation of predicted organokine effects in hiPSC-derived cardiomyocytes

To explore how the effect of the predicted adipose-to-heart signals may be evaluated we treated human induced pluripotent stem cell (hiPSC)-derived cardiomyocytes with recombinant human LGALS3 and THPO (**Figure 4A**). RNA sequencing of the treated cells shows significantly differentially regulated genes (DESeq2 s-value ≤ 0.1) compared to control cells (**Figure 4B**). Genes differentially regulated by galectin-3 treatment include those that are involved in predicted processes including the ribosome (*RPS29*, *RPS27*), as well as processes such as protein degradation (*MMP10*), immune process (*C4A*, *C4B*, *RELB*), and calcium signaling (*CAMK4*). By contrast, THPO treatment led to different sets of differentially expressed genes, including those involved in transcriptional regulation (*MED26*, *H4C16*), ribosome (*RPS27*), and ubiquitination (*UCHL1*) (**Figure 4C**). Gene set enrichment analysis (GSEA) of the RNA sequencing results corroborates significant up-regulation of ribosome genes upon LGALS3 (**Figure 4D–E**) and THPO (**Figure 4F–G**) treatment. Thus, treating hiPSC-derived cardiomyocytes with recombinant organokines at physiological concentrations present a viable strategy to experimentally corroborate the predicted ribosome associations, and may be useful for delineating the signaling consequences of other endocrine messengers.

**Figure 4.**
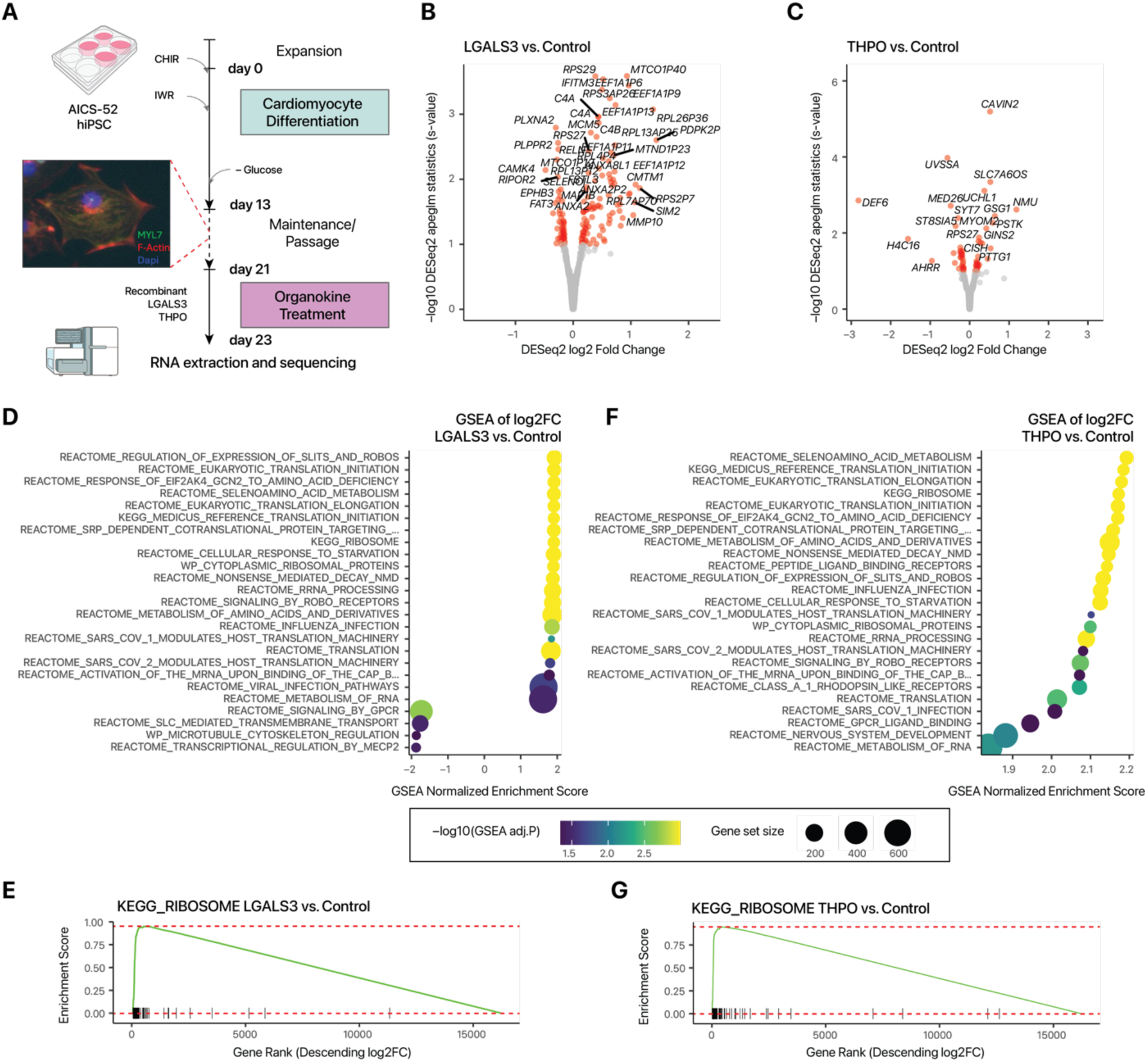
Validation in human induced pluripotent stem cell models. **A.** Experimental schematic of human induced pluripotent stem cell (hiPSC) differentiation into cardiomyocytes and recombinant organokine treatment. **B.** Volcano plot of RNA sequencing results showing significantly differentially expressed (red) (DESeq2 apeglm s-value ≤< 0.1) genes in hiPSC-cardiomyocytes treated with 14 ng/mL recombinant human LGALS3 proteins for 48 hours. **C.** As in B, but for hiPSC-cardiomyocytes treated with 0.1 ng/mL recombinant human THPO proteins for 48 hours. **D.** Dot plots of significant gene set enrichment analysis (GSEA) results showing the enrichment of ribosome terms I the gene expression profiles of hiPSC-cardiomyocytes treated with 14 ng/mL recombinant human LGALS3 proteins for 48 hours. **E.** GSEA enrichment plot of genes annotated as KEGG RIBOSOME in the RNA sequencing data, showing their preferential enrichment among up-regulated genes. **F.** As in E, but for hiPSC-cardiomyocytes treated with 0.1 ng/mL recombinant human THPO proteins for 48 hours. **G.** As in G, but for hiPSC-cardiomyocytes treated with 0.1 ng/mL recombinant human THPO proteins for 48 hours.

## Discussion

Recent years have seen rising interest in mapping the complex networks of interorgan signaling, such as the crosstalk of adipose tissues, liver, and the heart in cardiometabolic diseases (2). However, unraveling biologically meaningful crosstalk events remains a challenge due to the pleiotropy and ubiquitous expression of many circulating proteins. Here, we describe a computational approach to infer interorgan communication. This workflow builds upon and extends prior work to provide functional context of candidate signaling events, by considering the association between secretome-coding genes in the source tissue and transcriptome latent space representations in the target tissue. Decomposing the gene matrix into latent space representations using PLIER has the benefit of improving interpretability through prior knowledge annotations such as REACTOME and KEGG. These annotations are useful when assessing which processes were affected by secreted factors. For example, adiponectin along the adipose (subcutaneous) ∼ heart (left ventricle) axis is significantly correlated with processes involved in immunity and calcium signaling. The former process likely functions through resident immune cells that are intertwined with the heart muscle, whereas the latter is known to have a multitude of influences on both immune and cardiac cells such as contractility. The improved biological interpretation of the gene modules circumvents the complications regarding the inability to distinguish between cardiac and immune cells in the GTEx v8 data.

### Limitations of the study

As with other endocrine prediction strategies, SALVE is intended for the hypothesis generation purposes. Not every bona-fide endocrine signal may be expected to turn up as positive hits, as not every functional interaction proceeds through differential expression levels. For instance, we did not find well-established signals such as cardiac natriuretic peptides and adipose leptin, which could be due to their complex post-translational regulations (37) or non-linear concentration-dependent effect on tissues (1). In another instance, adiponectin is known to improve insulin sensitivity in vivo (38), but our co-expression analysis shows a negative correlation between adipose *ADIPOQ* and heart L.var.39 (KEGG Insulin Signaling Pathway; r: – 0.33, adj.P 3.7e–4) (**Supplemental Table S1**), ostensibly suggesting a desensitizing effect. This indicates the signs of population scale correlations (i.e., positive vs. negative correlation) may omit important biological contexts and should be interpreted with caution.

In parallel, we emphasize that candidate hits must be rigorously validated, as cross-tissue co-expression is correlational in nature and thus subject to confounding variables and reverse causality. Correlates including body weight may require further corrections to uncover true signals. This may also explain that in our analysis subcutaneous adipose tissues form more associations with other organs than visceral fat, even though the latter is thought to be more secretory. Other genetic and non-genetic variations can independently affect expression of two genes in two organs and give the appearance of interorgan signals, e.g., genetic variants that simultaneously influence the expression of two unrelated genes. Prior work has attempted to correct this with some success using a genetic de-correlation approach (15). Reverse causality may also occur in instances where genes in one tissue regulate circulating metabolites that then affect secretome expression in another organ in complex feedback networks. Prior work in transformed cell lines (e.g., C2C12 myotubes) has estimated that only 15–25% of predictions might be successfully replicable in experiments (14). However, it may be expected that the validation rate would be higher when physiologically more relevant models are employed for testing, including but not limited to the hiPSC-derived cardiomyocytes we used in this study. Likewise, because GTEx v8 provides extensive bulk rather than single-cell transcriptomics data for co-expression analysis, it is not clear from which cell types a secreted factor and pathway might originate; some of the effects *LGALS3* and *THPO* have on the heart may involve non-myocytes. Hence, we foresee the true validation rates of predictions will continue to improve in future work as next-generation microphysiological systems (e.g., hiPSC-derived organoids or engineered heart tissues) become increasingly used as experimental models.

Notwithstanding these limitations, we show that the approach is already useful for generating new insights. Applying this workflow to find new candidate signals between distal tissues and the ribosome pathways of the heart, we observe galectin-3 (*LGALS3*) and thrombopoietin (*THPO*) to be significantly associated with multiple cardiac pathways. Galectin-3 is known to be secreted from the adipose tissue from adipocytes and macrophages and is elevated in obesity (39). In the heart, galectin-3 modulates multitudinous processes spanning immune regulation and extracellular matrix remodeling (40). Our investigation corroborates significant association of galectin-3 on cytokine signaling in immune systems (**Figure 3B**), recapitulating its known roles in neutrophil and macrophage biology in the literature, as well as other cellular processes beyond immune response. Associated pathways involving ribosome, protein homeostasis (ubiquitin-mediated proteolysis), and energy metabolism (insulin signaling) indicate a broad influence of galectin-3 on cardiac function. In parallel, whereas the best characterized role of thrombopoietin is in platelet formation, it also has known physiological roles in the heart involving injury response to myocardial infarcts (41). Our predictions in turn suggest that circulating thrombopoietin, including from the liver pool, may regulate cardiac transcriptional and translational pathways (**Figure 3B**).

In summary, we describe a bioinformatics method using latent representations of massive transcriptomics data to infer endocrine signal candidates between different tissues. Considering ongoing challenges in understanding interorgan signaling, the presented approach may help prompt new research avenues and add to the armamentarium of contemporary methods to elucidate the downstream function of endocrine relationships. The results are in **Supplementary Table S1** and the code for generating new predictions from other data sources is freely available (see Data Availability).

## Data Availability

This study uses GTEx v8 data from the GTEx consortium. RNA sequencing data generated for this study are available on figshare at https://doi.org/10.6084/m9.figshare.29856230.v1. The code for SALVE is available on GitHub at: https://github.com/Lau-Lab/TissueCrossTalk_Prediction.git

## Acknowledgments

The authors thank members of the Edward Lau lab for helpful discussions.

## Disclaimers

The content is solely the responsibility of the authors and does not necessarily represent the official views of the National Institutes of Health.

## Author Contributions

M.P.L. and E.L. conceived and designed research; J.P. and P.S. performed experiments; J.P., C.K.V., and E.L. analyzed data; J.P., C.K.V., and E.L. prepared figures; J.P., C.K.V., M.P.L., and E.L. drafted, edited, and revised the manuscript. All authors approved the final version of the manuscript.

## Supplemental Figures

**Suppl. Figure S1.**
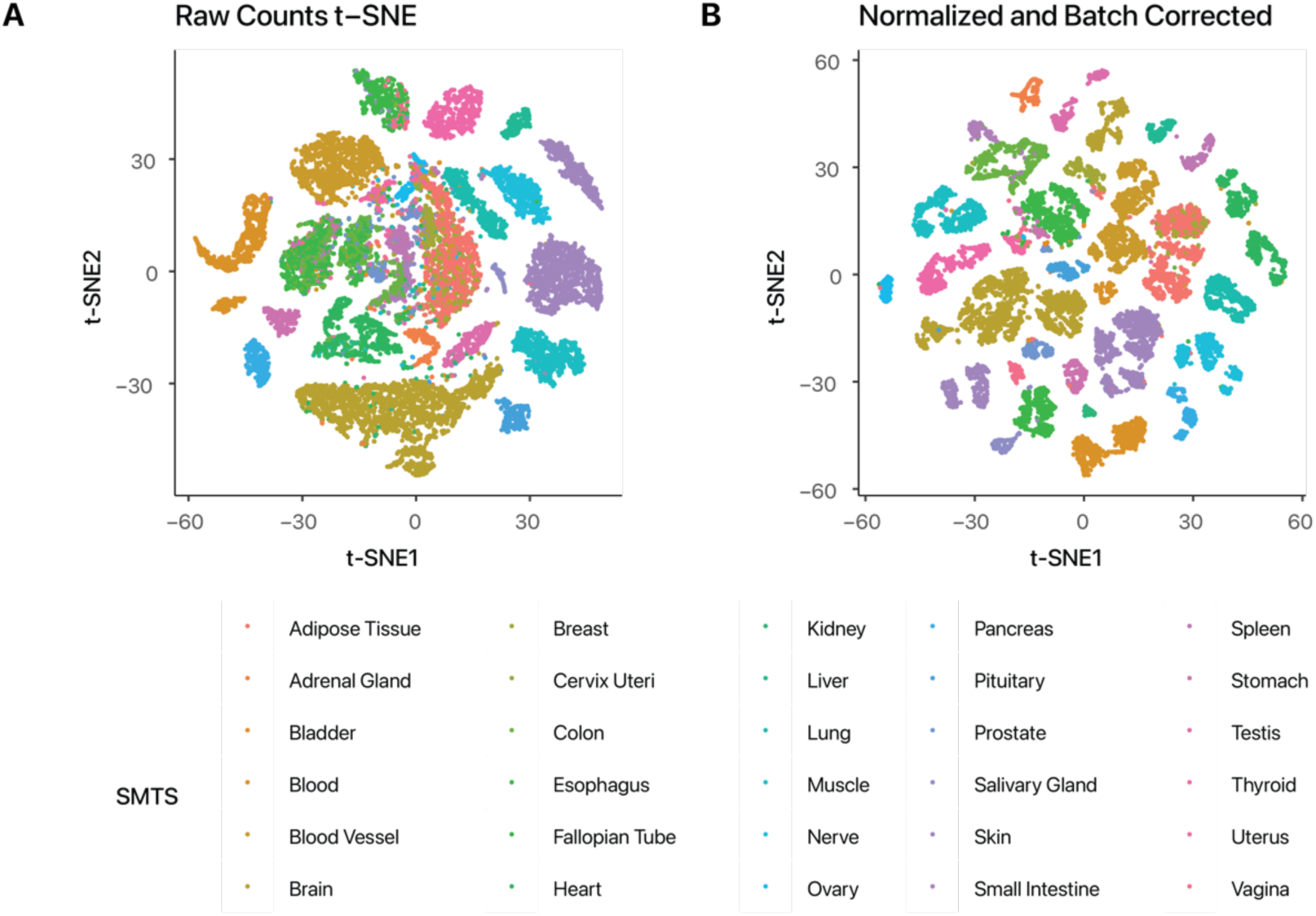
Tissue distributions of GTEx v8 transcriptome data. **A** Plot of t-distributed stochastic neighbor embedding (t-SNE) of GTEx v8 transcriptome data grouped by gross tissue type (SMTS). Tissue subtypes are not further distinguished by color. **B.** Plot of t-SNE of data following normalization and batch correction steps.

**Suppl. Figure S2.**
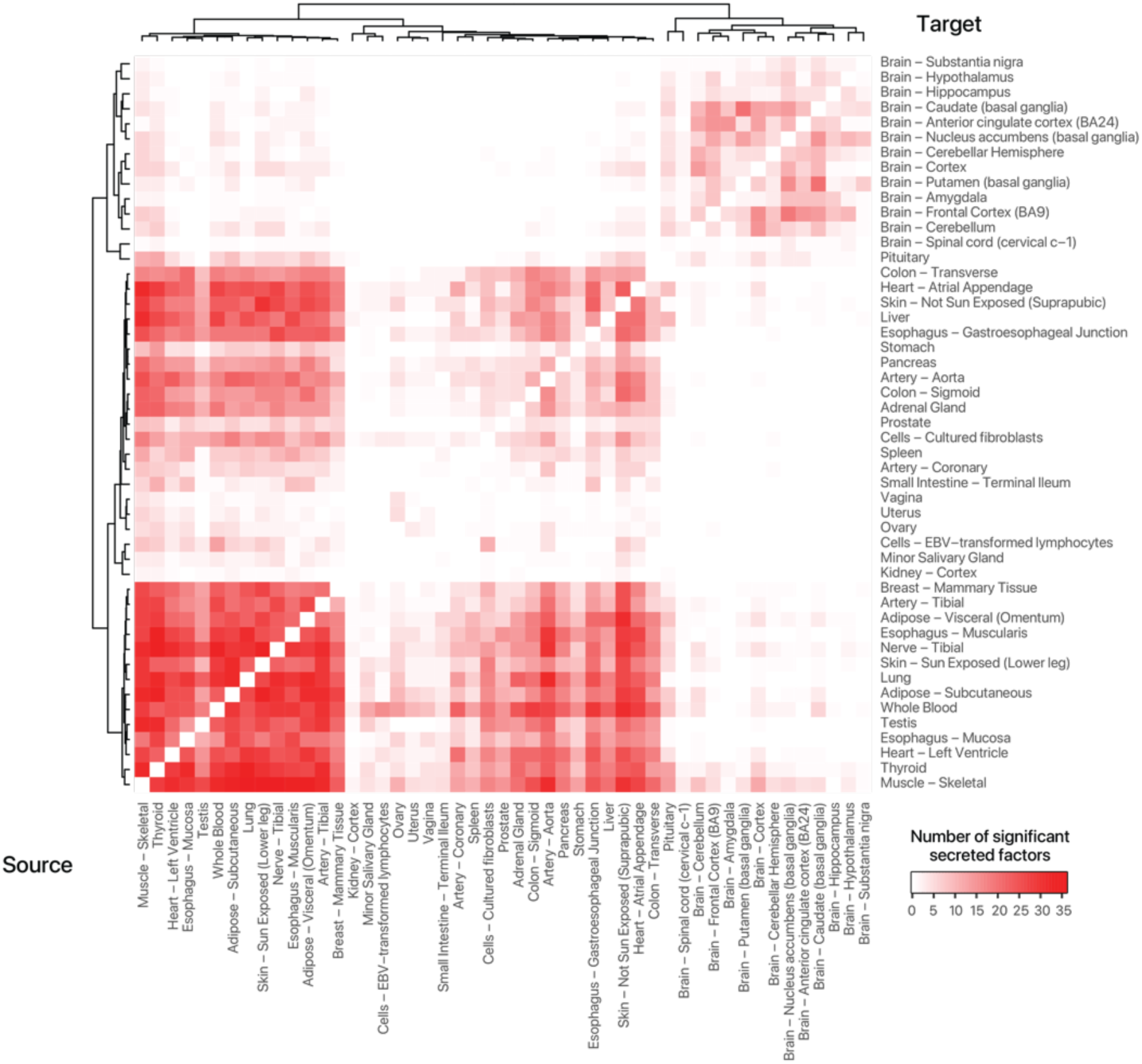
Significant gene to latent variable associations clustered by tissues. Heatmap showing significant correlation among prioritized organokines (adj.P ≤ 0.005, S* ≥ 2) across all source-target tissue pairs.

**Suppl. Figure S3.**
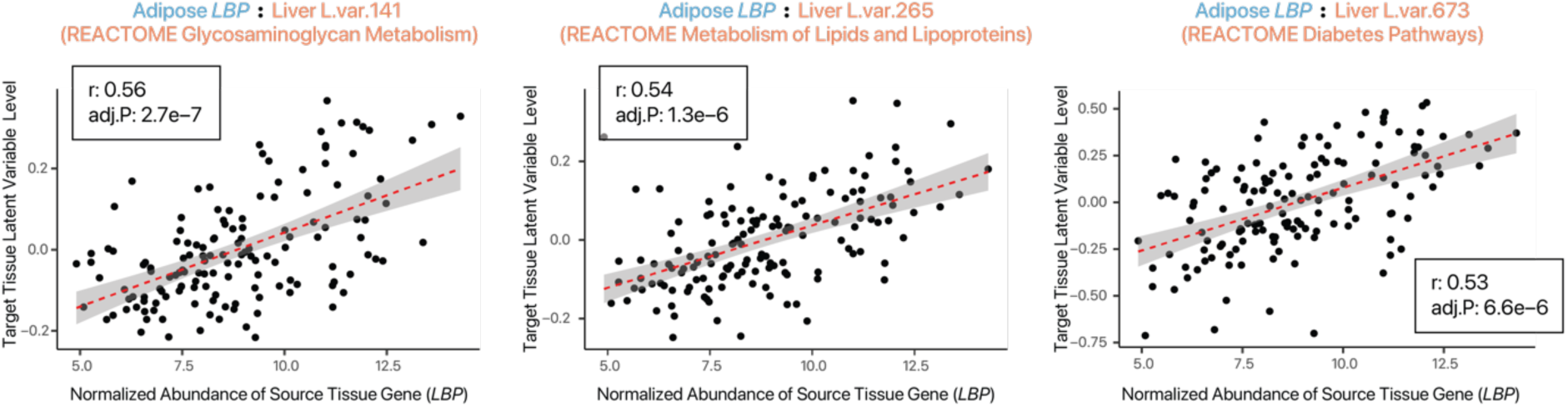
Association of adipose *LBP* with liver metabolism. Scatter plots showing the correlated expression of adipose (visceral – omentum) and liver latent variables related to glucose and lipid metabolism. Each data point is an individual donor sample with both adipose and heart transcriptomes in GTEx v8. Red dashed line: best fit linear fit. Ribbon: s.e.

**Suppl. Figure S4.**
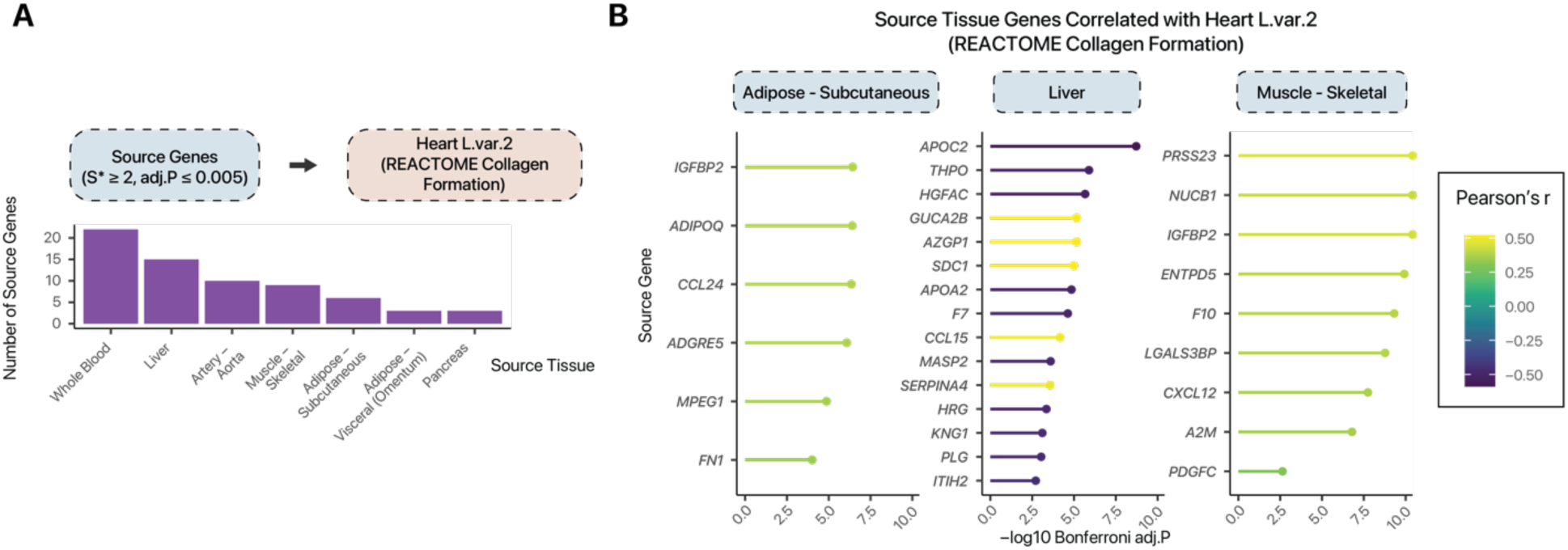
Candidate organokines associated with cardiac collagen formation. **A** Bar chart showing the number of significant organokines from several cardiometabolically relevant tissues that are significantly associated with heart latent variable 2 with REACTOME Collagen Formation annotation. **B.** Dot plot showing the correlation coefficient and significance of organokines from the adipose tissue (subcutaneous), liver, and skeletal muscle with heart latent variable 2 (REACTOME Collagen Formation).

